# Cryo-EM-based examinations of honey bee colony losses reveal structural insights into acute bee paralysis virus maturation and mechanical carriage

**DOI:** 10.64898/2026.11.22.768463

**Authors:** Jorge H. Medina-Duran, Judit J. Penzes

## Abstract

Honey bee colony losses pose a complex and significant agricultural problem in the United States and Europe, which has also been linked to viruses. As opposed to a purely sequence-based approach, we used cryogenic electron microscopy (cryo-EM) as an alternative sequencing- and culture-independent diagnostic tool to investigate viral pathogens associated with colony losses in the state of Texas. By purifying a limited number of viral particles directly from degraded hive remains, we could establish the identity of acute bee paralysis virus (ABPV) at the species level without prior sequencing data at 3.1 Å resolution, reconstructed of only 443 particles. We showed that small hive beetles carry mature ABPV virions, both within their digestive tract and attached to their cuticle. We successfully utilized cryo-EM to infer active ABPV replication from honey bee carcasses by reconstructing the virion structure of immature provirions at 2.35- and 2.53 Å, respectively, even several days after the collapse, despite the degradation of negative-strand RNA intermediates. Structural characterization of these immature particles revealed that ABPV capsids assemble around a condensed genomic core with the nucleic acid gradually expanding to occupy the capsid interior, ultimately bringing the VP0 structural protein in contact with VP3, which cleaves VP0 to structural proteins VP2 and 4. Together, our findings establish cryo-EM as a complementary diagnostic approach that simultaneously enables viral identification, replication-state assessment, and structural characterization, providing unprecedented insights into the maturation process of aparaviruses, a lineage of honey bee-infecting picorna-like viruses at the forefront of colony losses.

**Importance:** Metagenomics, as a diagnostic tool, yields exclusively sequence-based data, which offers limited insights into the complexity of viral diversity, abundance, biology and replication status. Cryo-EM is sufficiently matured to offer an alternative, sequence-independent, structure-first diagnostic approach to overcome these limitations, which we demonstrated by investigating honey bee colony losses that were accompanied by sever small hive beetle (SHB) infestations. The loss of honey bee colonies has reached an all-time high in the United States, with viral agents acting as prominent drivers of these losses. We showed that cryo-EM can establish species-specific viral identity from a limited number of particles, directly from highly degraded field-collected materteral, while distinguishing an ongoing viral infection from nucleic acid contaminations. This diagnostic strategy provided the first structural characterization of the ABPV virion and revealed that dicistoviruses undergo a fundamentally different maturation process from what was previously inferred based on other picorna-like honey bee viruses.

## Introduction

The field of detection, diagnosis, and identification of uncultivated viral agents has been dominated by metagenomics, revolutionizing diagnostic virology [1]. Despite being a relatively fast, high-throughput approach, metagenomics provides sequence-based information, with significant biases towards specific nucleic acid types, such as dsDNA [2]. Furthermore, distinguishing latent, episomal viral genomes or nucleic acid contaminants from true, replication-ready, mature virions poses limitations [3]. *De novo* short-read assembly may fail to accurately distinguish closely related viral genomes with high nucleic acid sequence identity, making them prone to misassembles [4]. These pitfalls can ultimately lead to false positives, misdiagnosis, and inaccurate estimates of true virion abundance, with limited insights into the biology of the viral pathogen.

The western European honey bee (*Apis mellifera*), is arguably one of the most valuable pollinators worldwide [5, 6]. Approximately, one third of all agriculturally significant plants depend on these insects for pollination [7]. Honey bee populations have been exhibiting historically high levels of colony losses in the United States [8, 9], recognized as a complex, multi-faceted issue of chemical stress, habitat loss, climate change, and pathogens [10–12]. Viruses are among the major drivers of these losses [12]. There are currently 24 viral agents associated with honey bees, out of which seven are linked to the disorder, all being +ssRNA viruses of the *Picornavirales* order [13, 14]. Within the *Picornavirales*, the *Dicistoviridae* includes four prominent honey bee viruses, namely acute bee paralysis virus (ABPV), Israeli acute paralysis virus (IAPV), Kashmir bee virus (KBV) and black queen cell virus (BQCV). ABPV, IAPV, and KBV are all classified as distinct species within the genus *Aparavirus*.

The invasive, ectoparasitic mite, *Varroa destructor,* is responsible for vectoring most of the honey bee viruses associated with declining colony fitness, including deformed wing virus (DWV), another picorna-like honey bee virus from the *Iflaviridae* family [15]. Little is known, however, concerning the role of the small hive beetle (SHB) (*Aethina tumida*) in vectoring honey bee viruses despite of it being an omnipresent hive pest [16, 17].

Several picorna-like honey bee viruses have been structurally characterized [18–21]. These studies led to an increased understanding of the genome release and uncoating process of aparaviruses and iflaviruses [18–20]. While the iflavirus DWV has been shown to assemble premature empty capsids to package the viral genome [18] the maturation process of aparaviruses remains uncharacterized.

Here, we used cryogenic electron microscopy (cryo-EM) and 3-dimensional image reconstruction to identify the viral agent associated with a colony collapse event in the southern US, accompanied by severe SHB infestation. We showed that cryo-EM is capable of sequencing-free identification of a member from the *Aparavirus* complex to the species level, even multiple days post loss, from a limited number of viral particles. This diagnostic approach yielded the first structural characterization of the ABPV virion. We show that SHBs carry intact ABPV virions in their mature state, which harbor surface-related conformational changes. We also used cryo-EM to assess whether ABPV was actively replicating in deceased honey bees prior to its loss. Lastly, our diagnostic pipeline also provided the first structural reconstruction of the ABPV virion at three maturation stages, shedding light on the aparavirus maturation process.

## Results

### Cryo-EM-driven virus identification

In July 2025, a honey bee colony in Bastrop, TX, collapsed, leaving only a small number of deceased honey bees, while larval SHBs were in abundance. The destroyed frames were collected and transported to our laboratory five days later, as an assemblage of fragmented comb, SHB larvae, and scarce honey bee carcasses (Fig. 1A). As the remains lacked signs of fungal- or bacterial growth, a viral agent was suspected. We subjected 100g of fragmented beeswax, mixed with SHB larvae, to virus purification on sucrose cushion and gradient. Faint blue scattering was observed in the 25-50% fractions, which were pulled. Following high-resolution cryo-EM data collection, scarce isometric particles of ∼30 nm in diameter were observed in the micrographs (Fig. 1B, data collection parameters and refinement statistics included in Table S1, local resolution map in Fig. S1). The micrographs were clean and mostly free from protein debris, bacteriophages, or other contaminating agents. Despite boxing only 443 particles, we could still reconstruct the 3-dimensional structure of a virion of T=p3 icosahedral symmetry at 3.1 Å resolution (Fig. 1C, Fig. S1). Although the 2-dimensional averages of the particles indicated the presence of a uniformly electrodense interior (Fig. 1B), the density was weakly ordered in the particle center (Fig. 1C). The electron density map was subjected to automated *de novo* model building, using the ModelAngelo software[22], without providing any sequence data to guide the identification of residues within the electron density (Fig. 1D). 180 contiguous modeled residues were subjected to homology search using the BLASTp suite (Table S2). The lowest E-value hit of 1.65E-69, was to the NCBI cluster of ABPV, specifically to structural protein (VP) 2, distinguishing the 180-aa-long sequence from those of IAPV (E-value of 4.3E-69) and KBV (E-value of 4.78E-69). This species level assessment was further supported by slight conformational divergence in the modeled Cα backbone compared to IAPV or KBV, as well as by density shape differences in eight single residue positions, which differ in the ABPV VP2 compared to at least one of its close relatives (Fig. 1E).

**Figure 1.**
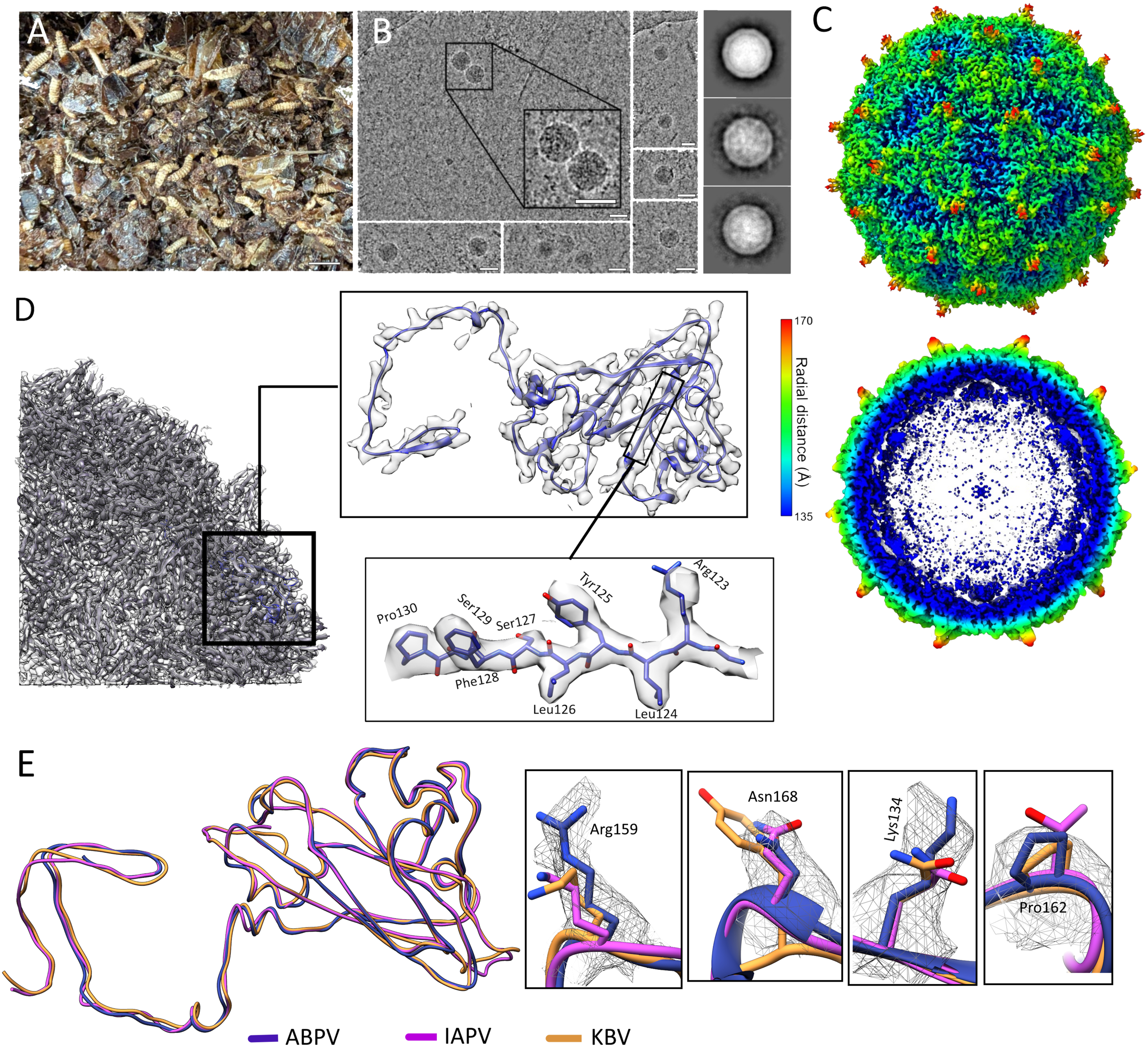
Sequencing-free, Cryo-electron microscopy (cryo-EM)-mediated virus identification in a collapsed honey bee colony. (A) Representative image of the diagnostic material our laboratory received for diagnostic purposes, including fragmented comb, small hive beetle larvae, and scarce honey bee carcasses. (B) Cryo-EM micrographs showing isometric particles of 30 nm in diameter, harboring an electrodense center. The scale bars indicate 30 nm. (C) Radially colored electron density map of the isometric particles, presented in the i,222 icosahedral orientation. The left panel shows the surface, while the right panel displays a cross section of the particle. The electron density map was reconstructed at 3.1 Å resolution. (D) Identification of the recovered viral particles by *de novo* model building directly into the electron density map at 3.1 Å resolution. A subsection of the density volume was isolated and modeled. The left panel shows the density section with the *de novo* atomic model as ribbon diagrams. The longest and least fragmented model chain was extracted, as shown by the top insert. The bottom insert displays an example of the automatically built *de novo* atomic model, with residue numbers reflecting their position in this extracted chain. (E) Superimposition of the *de novo*-built chain backbone of acute bee paralysis virus (ABPV) with the VP2 backbone models of Kashmir bee virus (KBV) and Israeli acute paralysis virus (IAPV). The right insert shows example residues, which were correctly identified in ABPV and represent single polymorphic residues compared to IAPV and KBV. The electron density of the 3.1 Å ABPV map is shown as a mesh, while the IAPV and KBV models were fitted into this density for comparison.

### Genome sequencing and atomic model refinement

As ABPV is known to harbor a ssRNA genome, we specifically targeted this nucleic acid type for sequencing efforts to corroborate the structural ID. Thus, reverse-transcribed RNA of purified virions was subjected to short-read Illumina sequencing. The near-complete viral genome encompassed 9,637 nucleotides (nt), missing only ∼70 nt at the 3’ end. The genome organization was consistent with that of dicistoviruses, confirming the ABPV identity of both the non-structural and structural polyproteins (95.29% and 94.81%, respectively). The 90.54% nt-level identity to the reference genome designated the sequence as a new genotype.

The *de novo* atomic model provided guidance to assign each sequenced VP to the electron density (Fig. 2A). This process successfully identified VPs 1, 2, and 3 (Fig. 2B), with their ordered residue range summarized by Fig. S2. To model VP4, we inspected the capsid interior manually and located the corresponding density at a slightly lower σ contour threshold. VP4 was ordered in its entire length except for its first 16 N-terminal residues. The jelly roll cores of three major VPs were ordered even at high contour threshold values (∼4 σ) (Fig. 2C). Our structural data corroborated with the previously proposed protease cleavage sites [23, 24] (Fig. S2).

**Figure 2.**
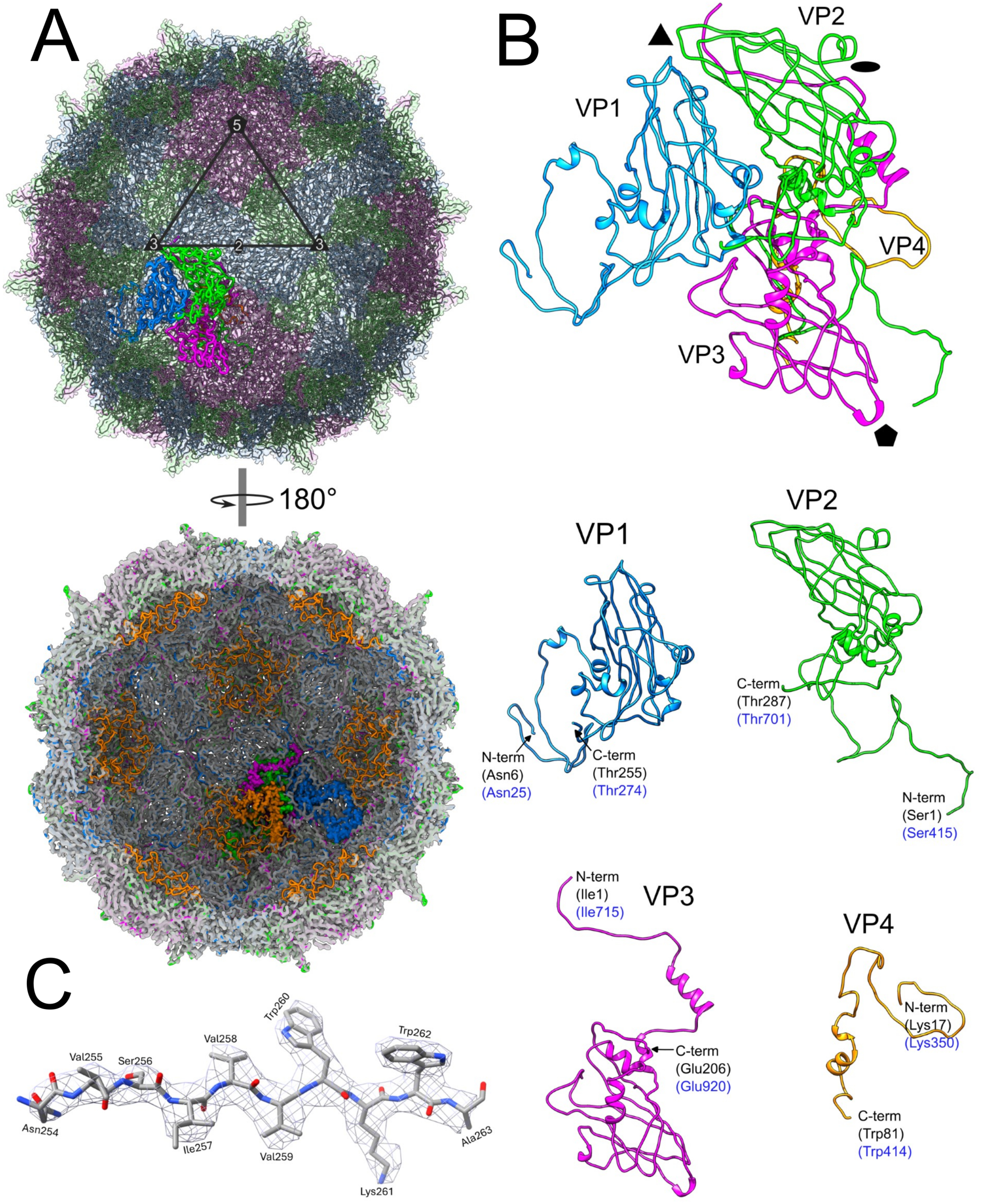
Structural characterization of the acute bee paralysis virus (ABPV) virion structure. (A) Surface (top) and interior (bottom) view of the resolved electron density map, showing the atomic model, which was constructed based on the genome sequencing results. Capsid proteins (VPs) are shown in various ribbon colors, with the triangle highlighting an asymmetric unit (ASU) of the four VPs. The fivefold symmetry axis is marked by a pentagon, while the triangle indicates the threefold axis, and the ellipsoid the twofold axis. VPs comprising an ASU are highlighted by the thick and vibrantly colored ribbon diagrams. (B) Ribbon diagrams of a single ASU, shown in the colors corresponding with those of panel A. The bottom panel displays the ribbon diagrams of each VP separately. (C) Example density of the modeled electron density map. The electron density is shown as a grey mesh, occupied by the refined atomic model.

### The ABPV virion structure in a *Dicistoviridae* context

The ABPV virion displayed a similar surface morphology to the honey-bee-infecting members of the *Aparavirus* genus (Fig. 3A). Phylogenetic inference of the aparaviral VPs indicated that ABPV is possibly evolutionary equally distant from KBV and IAPV (Fig. 3B). Superimposition of the ABPV VP atomic models with those of IAPV and KBV showed low RMSD values overall. The lowest pairwise RMSD values were calculated between KBV and IAPV, supporting their closer structural similarity. The KBV VP4, however, was marginally more similar to ABPV (RMSD = 0.701) (Fig. 3B). Deviations in the otherwise remarkably similar backbone structures were restricted to flexible regions, i.e., the surface spikes, the fivefold- and the threefold symmetry axis, and VP4. In case of VP4, seven additional residues could be N-terminally modeled in comparison to the IAPV and KBV structures.

**Figure 3.**
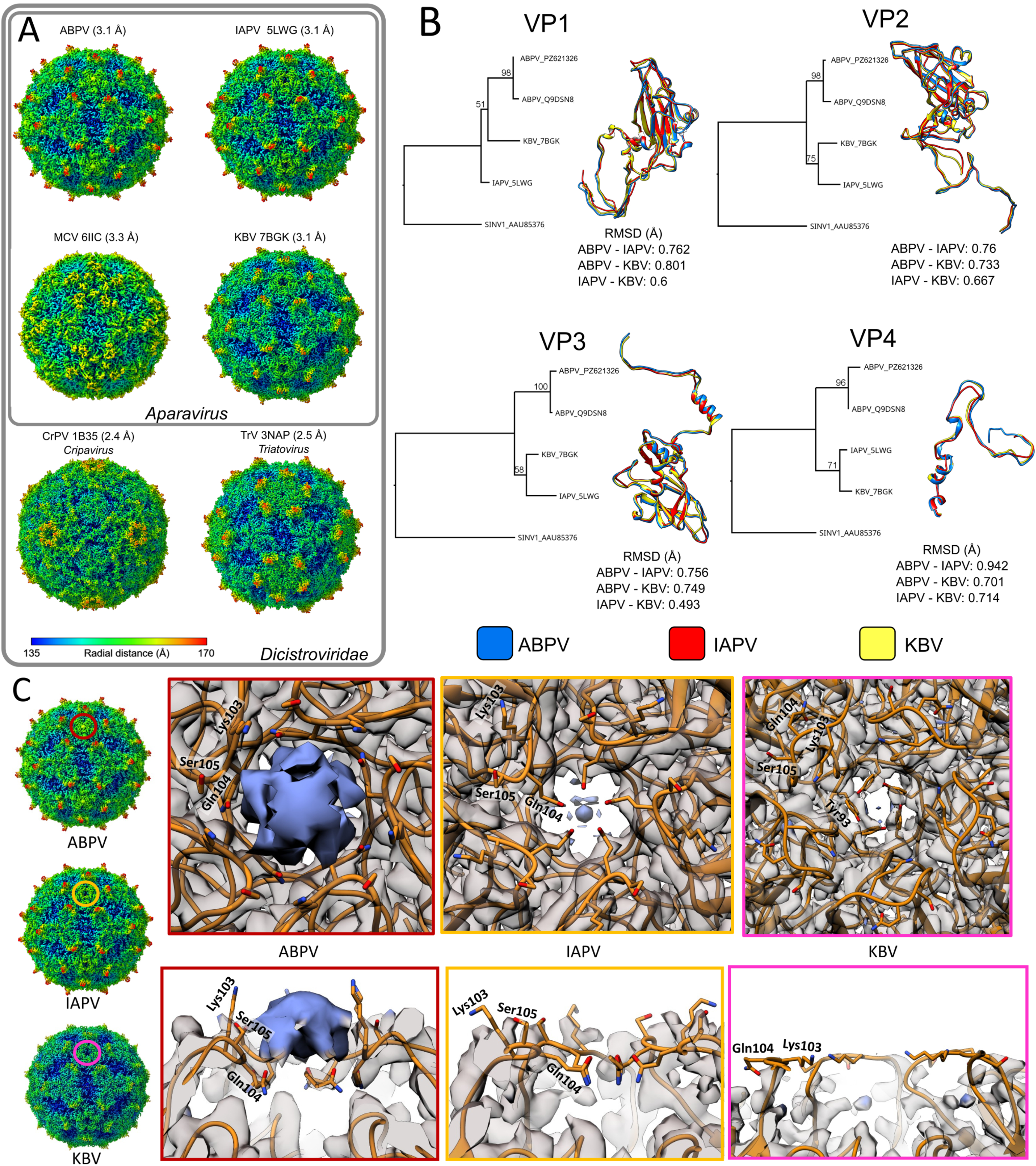
Acute bee paralysis virus (ABPV) in the structural context of the *Discistoviridae* family and the *Aparavirus* genus. (A) Surface comparison of the ABPV virion structure to those of other dicistroviruses. Note the prominent surface spikes, which are characteristic of the honey bee-infecting members of the *Aparavirus* genus. The density maps are oriented in the i,222 icosahedral convention. Abbreviations: IAPV – Israeli acute paralysis virus, MCV – mud crab virus, KBV – Kashmir bee virus, CrPV – cricket paralysis virus, TrV – Triatoma virus. (B) Phylogenetic reconstructions of the relationships between each aparavirus capsid protein (VP). The ribbon diagrams show the results of the superimposition of the respective VPs with the obtained root mean square deviation values in ångströms (Å). (C) Comparison of the fivefold symmetry axis of the virion structures within genus *Aparavirus*, which is highlighted by the colored circles in the small electron density maps. The magnified images show the electron density map as transparent grey, if it could be modeled, while unmodeled densities are shown in blue. The top panels show the fivefold axis in top-down view, while the bottom inserts display the cross-section view. The atomic model is shown in yellow, and the residues coordinating the extra fivefold-related density in ABPV are labeled.

The fivefold axis opens in a pore in IAPV and KBPV, while a piece of unmodeled density covered this pore in ABPV, reaching ∼11 Å in diameter (Fig. 3C). Its presence alters the conformation of Lys103, Gln104 and Ser105, which exhibit an ordered backbone. The corresponding region is significantly more flexible in IAPV and KBV, with the three residues lacking backbone ordering.

### Limit of detection

We developed diagnostic PCR- and qRT-PCR-based systems, targeting the structural polyprotein gene, to establish the detection limit of cryo-EM in relation to PCR-based methods. We prepared a 10-fold dilution series, using the concentrated virus suspension from the cryo-EM workflow. Viral titers showed a nearly log-linear decline from the undiluted suspension to the 1:100 dilution, which had a titer of 10⁹ gc/mL (Fig. 4A). Faint bands still appeared at a concentration of 10⁷ gc/mL, while the qRT-PCR only produced sporadic amplification at high C_q_ values. Accordingly, the detection limit of ABPV RNA is ∼10⁷ gc/mL, with increased confidence at viral titers of 10⁸ gc/mL or higher.

**Figure 4.**
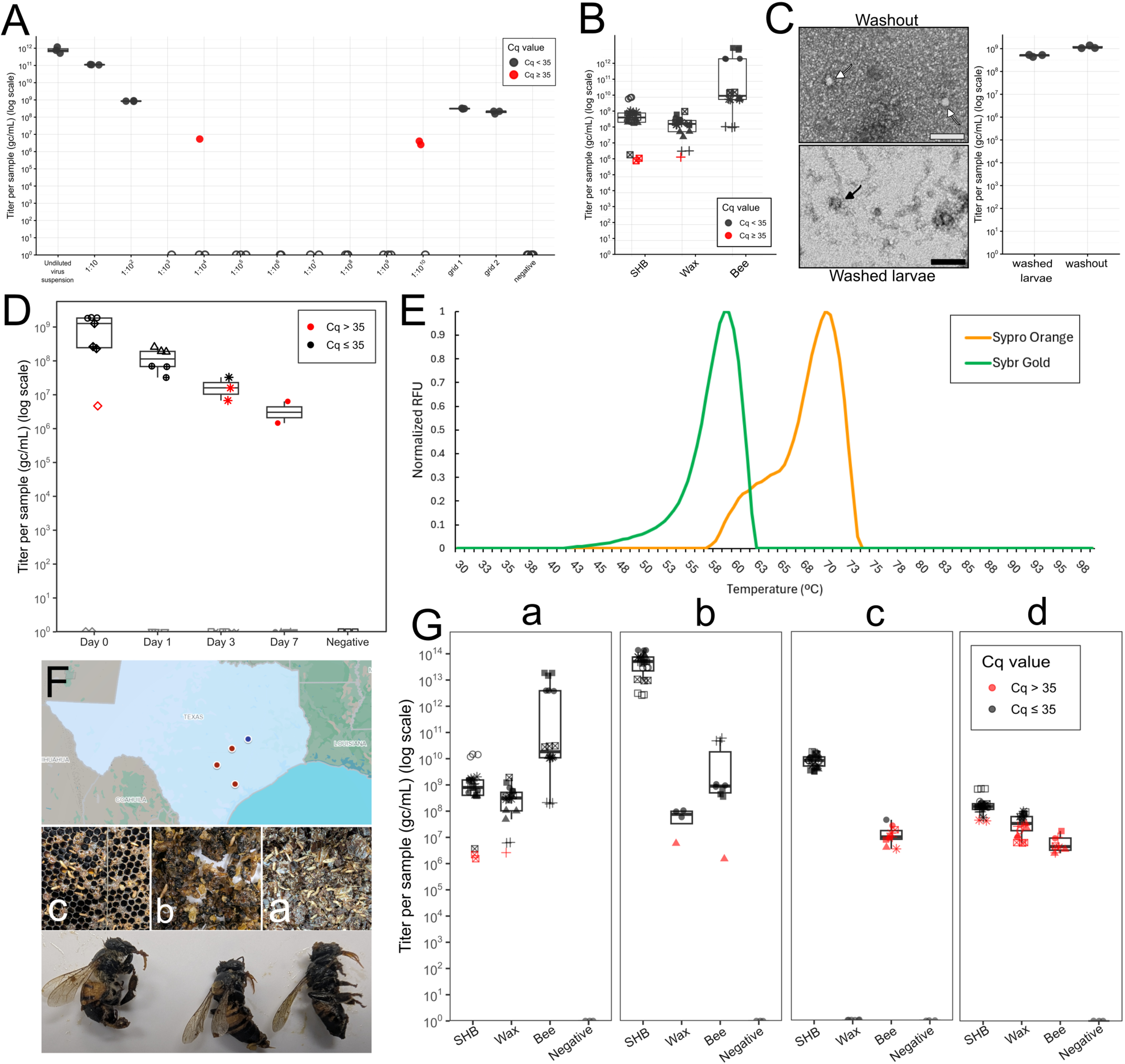
Assessment of acute bee paralysis virus (ABPV) viral titers in the context of detection limits, environmental stability, and multiple collapsed colonies. (A) Quantitative reverse transcription (qRT) PCR-determined viral titers of the 10-fold dilution series investigating the detection limits of the PCR-based ABPV detection method. The labels grid1 and grid2 indicate the replicates of ABPV quantification on the surface of blotted cryo-EM grids. (B) ABPV titers in the three types of diagnostic material from the collapsed Colony A, shown as box-and-whiskers plots. (C) Negative staining electron micrographs showing ABPV particles washed off from the cuticle of small hive beetle larvae and particles imaged in homogenized small hive beetle larvae. The scale bar measures 100 nm. The box-and-whiskers plot presents the viral titers obtained from the larval cuticle and from homogenizing small hive beetle larvae. (D) qRT-PCR-quantified titers of beeswax incubated at ∼30°C for the span of a week. (E) Differential scanning fluorometry, assessing the temperature of ABPV genome ejection (SYBR^TM^gold) and particle disintegration (SYPRO^TM^orange). (F) The locations of collapsed colonies A, B and C are shown as red dots in the top panel, mapped to the state of Texas. The blue dot indicates the location of the healthy Colony D. The middle panel shows examples of the diagnostic materials from colonies C, B, and A. The bottom panel presents honey bee carcasses from the collapsed Colony B, which all exhibit paralytic signs, manifesting as the extended proboscis. (G) Box-and-whiskers plots presenting the ABPV titer quantification results across the four sampled colonies.

Two cryo-EM grids, identical to the ones used to structurally identify ABPV, were blotted, washed and incubated in phosphate-buffered saline (PBS), overnight. After correcting for the dilution factor, the PBS wash exhibited a titer of 10⁸–10⁹ gc/mL (Fig. 4A). This suggests that cryo-EM can detect and structurally characterize viruses at on-grid concentrations that overlap with the reliable detection range of the conventional, PCR-based diagnostic methods. This was achieved by 100-fold concentrating a virus suspension in the 10^10^ gc/mL range.

### ABPV abundance and distribution within the collapsed colony

The identification of a viral agent associated with the collapsed hive prompted us to further characterize the role of ABPV in the loss. From the colony remains designated as Colony A, we extracted beeswax, SHB larvae, and honey bee carcasses. All three sample types yielded a positive result, with the highest titer of 10^12^ gc/mL exhibited by the bee carcasses (Fig. 4B). The SHB larvae and the beeswax also contained viral RNA, ranging 10^6^–10¹⁰ gc/mL. To locate the source of the SHB-related ABPV particles, the larvae were washed in PBS, and subsequently homogenized. Negative staining transmission electron microscopy (negTEM) visualized particles consistent in size and appearance with the ABPV virions in both the wash and the homogenate. The dilution-corrected viral yield indicated similar quantities of viral RNA in the homogenized larvae (5×10^8^ gc/mL) to the amount of viral genomes washed off the SHB cuticle (1.2×10^9^ gc/mL) (Fig. 4C).

### Environmental stability of ABPV virions

Beeswax samples were incubated at 30°C for seven days with daily sampling and subsequent RNA extraction. Viral RNA was detected at high titers at day 0 in two of the three tested biological replicates, with samples showing values ∼10⁸-10⁹ gc/mL (Fig. 4D). By day 1, viral titers had declined by approx. one order of magnitude, with only two of the three biological replicates testing positive. By day 3 only one biological replicate tested positive for ABPV with viral loads close to the detection limit of 10⁷ gc/mL, with only two positive technical replicates (10⁷ gc/mL) by day 7 (Fig. 4D).

ABPV virions were subjected to differential scanning fluorometry (DSF) in the presence of SYBR^TM^Gold nucleic acid- and SYPRO^TM^Orange protein stain at pH 7.4, to assess their thermostability (Fig. 4E). The ABPV genome is ejected at approx. 59 °C, with the complete particle disassembling only at 70 °C. This is identical to the genome ejection temperature of KBV [19] albeit lower than that of IAPV (63°C) [20].

### ABPV and colony losses in Southern Texas

We reached out to the Texas Apiary Inspection Service (TAIS) to locate lost colonies with severe SHB infestation. Within the span of a week, we obtained the remains of two more colonies (Fig. 4F). Colony B exhibited mass mortality of honey bees with signs of paralysis, including extended, unretractable proboscis. The remains of Colony C only contained comb and SHBs, with a history identical to Colony A. By the Spring of 2026, this apiary replaced the collapsed Colony C, from which we could obtain healthy honey bee samples. We also gained access to a healthy colony near Bryan, Texas, designated colony D. ABPV was detected in all three colonies, with the healthy Colony D exhibiting the lowest viral titers (Fig. 4G). As we could not collect SHB larvae from this colony, we only sampled adult beetles, which possessed the highest ABPV titer of 1.8×10^8^ gc/mL, compared to 4.3×10^7^ gc/mL of wax and 6.7×10^6^ gc/mL of healthy honey bees. Colony C exhibited a relatively high titer of 8.8×10^9^ gc/mL in the SHB larvae collected at the time of its collapse. We could not detect ABPV presence in beeswax from Colony C. Bees of the restored colony exhibited near-detection limit titers. Colony B displayed the highest viral titer in its larval SHB population (∼10^14^ gc/mL) while high viral titers were detected in the paralyzed honey bee carcasses (10^11^ gc/mL), as well as in the beeswax (6.6×10^7^ gc/mL).

### Cryo-EM-mediated examinations of viral replication

Active replication of +ssRNA viruses is evaluated by detecting and quantifying the negative RNA strand[25]. To establish whether ABPV caused the paralytic disease in Colony B the facts that all individuals were dead and had been stored at temperatures exceeding 30°C in transit, posed major obstacles, due to the rapid degradation of unprotected -ssRNA. To overcome this, we subjected 20 honey bee carcasses to the aforementioned virus purification protocol, but this time we only collected the fractions spanning 25-40%, aiming to only collect immature viral particles. Although the micrographs were abundant in various protein debris, we could still box 1221 particles identical to those collected from Colony A (Fig. S3A and B). 3-dimensional classification at 6 Å distinguished two classes of 832 and 312 particles, which were refined separately to two structures at 2.35 Å and 2.53 Å, respectively (Fig. S3C, local resolution maps shown in Fig. S4). Although both reconstructions exhibited an almost identical backbone conformation to that of the Colony A structure (Fig. S3D), there were considerable differences in ordered residue ranges (Table S3), as well as in the cleavage state of VP4 (Fig. 5). Both structures exhibited the fivefold-related unidentified density, which slightly differed in shape (Fig. S5).

**Figure 5.**
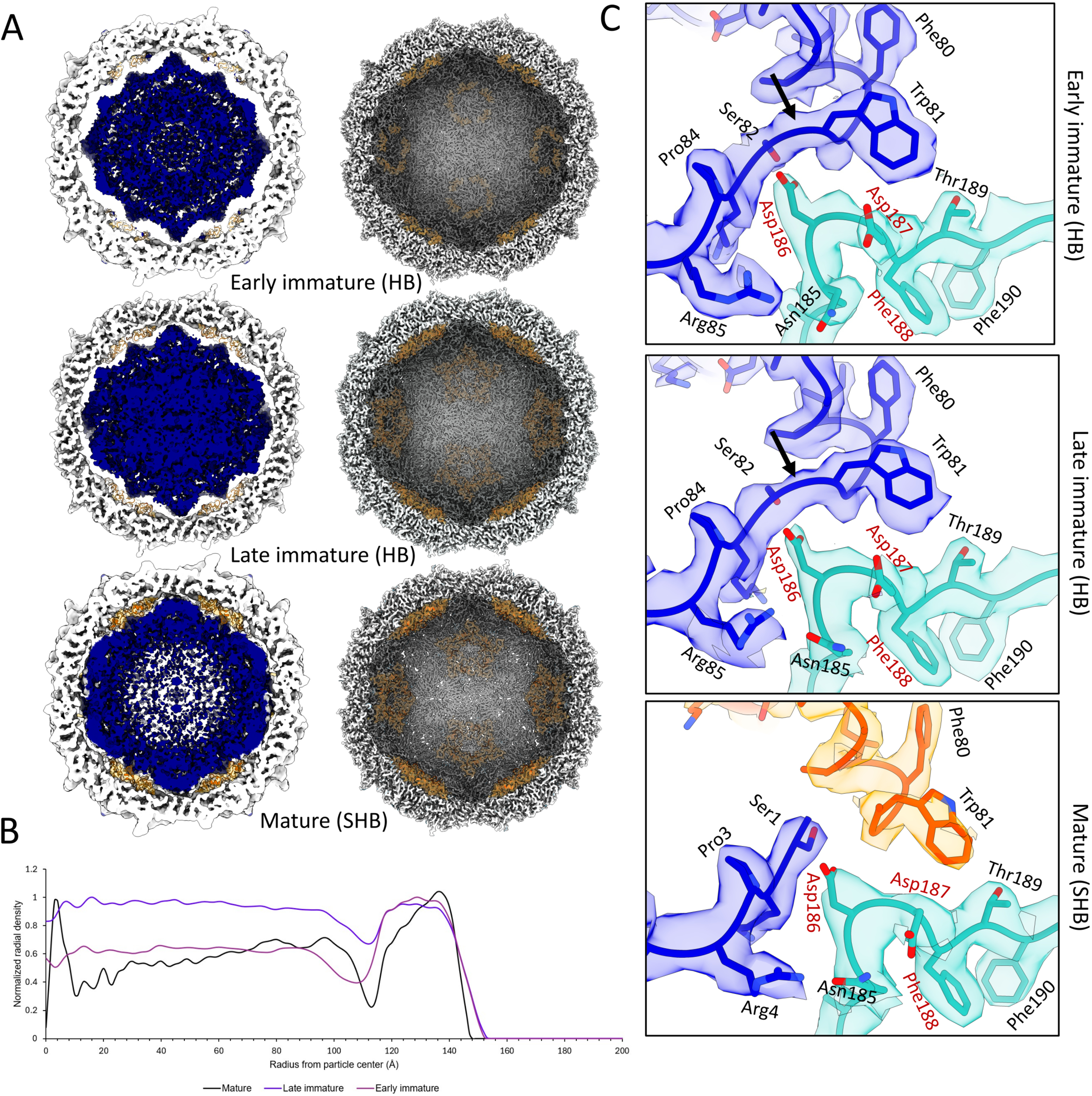
Structural characterization of the acute bee paralysis virus (ABPV) maturation process. (A) cross sections (left panels) and interior views (right panels) of three ABPV maturation stages. The ABPV capsid is shown in grey, while the density attributed to the ssRNA genome is presented in blue. The ordered residues of the structural protein (VP) 4 or the ordered N-terminal residues of VP0 are shown as orange ribbon diagrams, surrounded by transparent yellow electron density, with the genome-related density removed for presentation purposes. (B) Radial density distribution plot showing the amount of electron density measured at each Å from the particle center. (C) Density and atomic model corresponding to the residues, which play a role in the VP0 cleavage to VP2 and VP4. The black arrow indicates the cleavage site in the uncleaved VP0, shown as blue atoms in the blue density. VP3 is shown in teal with red labeling highlighting the residues of the DDF motif, hypothesized to perform the autocatalytic cleavage. The cleaved, matured VP4 is shown as an orange model in orange density.

### Structural characterization of ABPV virion maturation

The distribution of the luminal density, attributed to the ssRNA genome, differed considerably throughout the three reconstructions. The first honey bee-related class exhibited compact luminal density (Fig. 5A). A significant gap of ∼20 Å could be observed between the genomic density and the interior wall of the capsid, which was confirmed by its radial density distribution plot (Fig. 5B). This gap was also present in the second honey bee-derived class, measuring only 10-15 Å in width, and was accompanied by similarly well-ordered RNA density. The virion of colony A only exhibited a narrow gap of 4-8 Å, while the ordering of the RNA density gradually decreased towards the particle center. In contrast, VP4 was almost completely disordered in the structure with the largest density gap (18 ordered residues), while the number of the ordered residues increased as the width of the gap decreased, adopting first an additional 35 in the second class, then another 11 residues in the colony A population (Fig. 5A, Table S3). VP4 was still attached to the N-terminus of VP2 in both honey bee-derived reconstructions, comprising VP0, which confirms their immature state (Fig. 5C). The first population likely represents an earlier immature form of the ABPV virion, whereas the second class likely corresponds to a later immature form, as supported by the gradual increase of the VP4(VP0) density and the expansion of the viral genome, subsequently losing its intrinsic ordering.

At the VP4-VP2 cleavage site of VP0, Asp186 of VP3 comes in contact with Ser82 of the immature VP0, which then corresponds to Ser1 of VP4 following cleavage (Fig. 5C). This supports the IAPV-based hypothesis [21] that the DDF motif of Asp186, Asp187, and Phe188 of VP3 are responsible for the autocatalytic cleavage of VP0 in aparaviruses. In the mature particle, the 13 C-terminal residues of VP2 are absent from the virion structure, leaving a surface-exposed C-terminus at the base of the spikes (Fig.6). In both immature populations the corresponding terminus is completely ordered and flexible, adopting higher B-factor values as the maturation progresses (Fig. 6A and B). In the absence of this C-terminal segment, the surface spikes adopt more flexibility in the mature virions, while only their peak exhibits high B-factor in the immature virions (Fig. 6A). Similarly to IAPV and KBV, the C-termini occupy the canyon between the VP3 subunits, which constitute the fivefold axis (Fig. 6B). The flexible C-terminus is held in place by its second to last residue, Met380 (VP0 numbering), which establishes stacking interactions with the surrounding tyrosine-, arginine-, and glutamic acid side chains (Fig. 6C).

**Figure 6.**
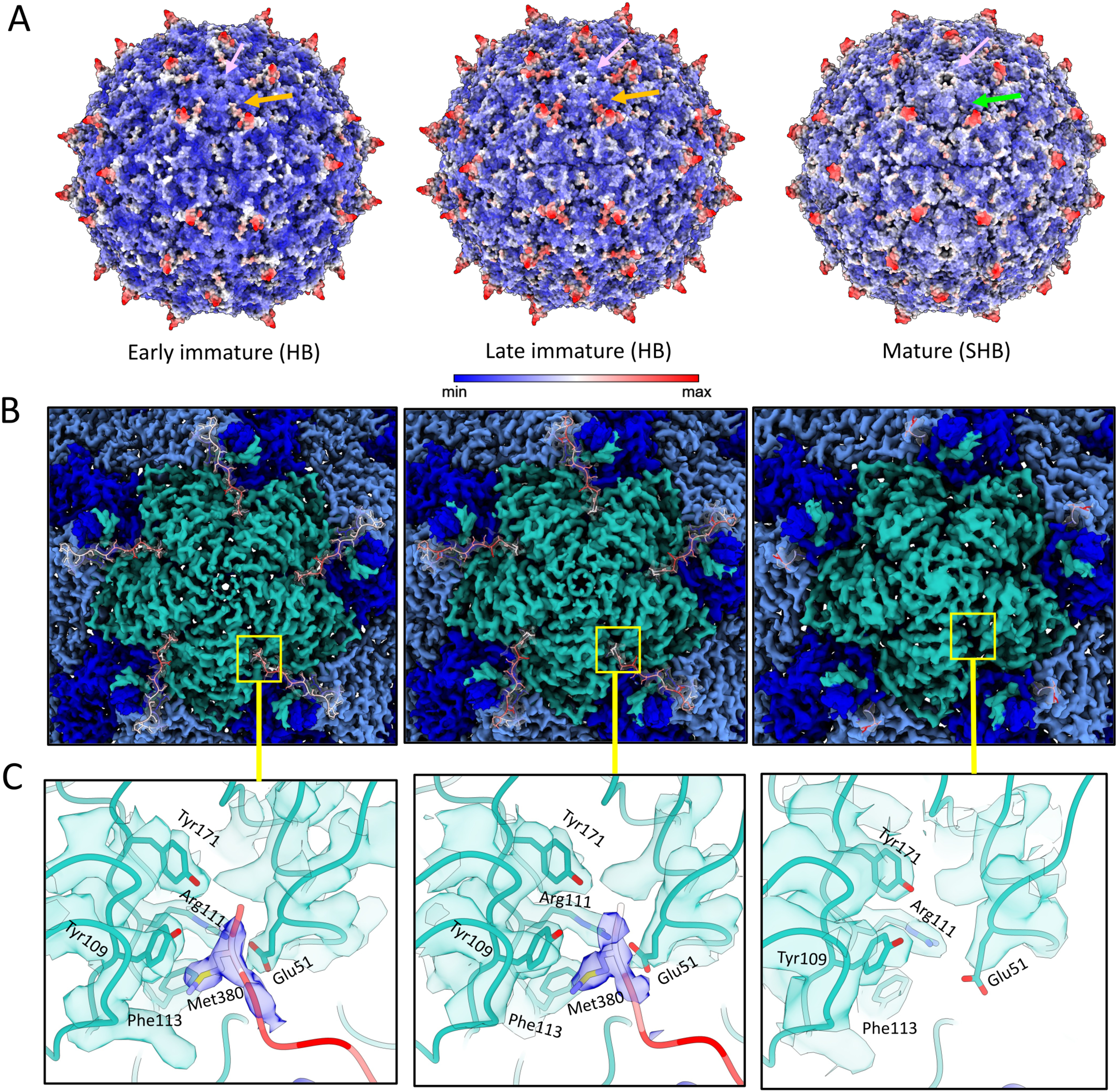
Structural analysis of the flexible C-termini of structural protein (VP) 2 and VP0 of the acute bee paralysis (ABPV) virion. (A) B-factor maps of the ABPV capsid surface in its three maturation stages. The yellow arrows indicate the ordered VP0 C-termini, while the green arrow points out the absence of the corresponding VP2 C-termini. The pink arrow indicates the fivefold symmetry axis. (B) Top-down view of the ABPV fivefold symmetry axis. The electron density map around VP3 is colored in teal, while the map is presented in light blue around VP1 and dark blue around VP2 or VP0, depending on maturation state. The VP2 or VP0 C-termini are shown as atomic models in transparent grey density, colored by their B-factor values. (C) Inserts showing the atomic models occupying the density at the fivefold canyon, which keeps the VP0 C-termini attached to the capsid surface. The electron density is zoned around the residues, which are involved in the interaction.

## Discussion

Here, we aimed to evaluate the suitability of cryo-EM as a diagnostic platform capable of providing species-level pathogen identification and insights into viral replication status directly from limited quantities of severely degraded, field-collected material. Although the potential of cryo-EM-mediated diagnostics has been previously documented [26], earlier work focused on identifying and characterizing the causative viral agent of a major invertebrate epidemic, for which abundant, freshly diseased specimens were available, enabling the recovery of high-quality viral material. In contrast, routine agricultural diagnostic investigations frequently rely on limited, poorly preserved specimens collected after the onset of disease or mortality.

Metagenomics and PCR-based viral detection are limited to sequence-based information, which only provides a restricted view of the complexity of viral outbreaks. We utilized a cryo-EM mediated, structure-first approach to investigate a case of honey bee colony loss, a complex and prominent agricultural and ecological problem [11, 27, 28]. By using an imaging technique, the diversity and abundance of encapsidated viral particles could be assessed immediately. From virus purification through high-resolution data collection to 3-dimensional reconstruction, the identity of the viral agent present in the collapsed colony was established within a week. Just by utilizing common grade EM grids, a general data collection pipeline, and only a single day of collection time, we obtained a 3.1-Å ABPV virion structure of only 443 particles. This resolution was sufficient to identify the viral agent at the species level, despite it being the member of a species complex of three closely related viruses [29]. This ID was confirmed by a traditional metagenomic approach, where we could specifically target ssRNA, as the genome type was already revealed by the sequence-independent identification. Concentration of the initial viral suspension (∼10¹⁰ vg/mL) enabled cryo-EM detection at an estimated on-grid equivalent of ∼10⁹ vg/mL, demonstrating that structure-based identification remains feasible at relatively low particle abundance.

Die-offs in farmed animals or plants in a typical agricultural setting may only be discovered hours or days following their onset. Diagnostic laboratories frequently rely on samples obtained from the public, which may not be preserved appropriately for subsequent diagnostic purposes [30–33]. In case of +ssRNA viruses the confirmation of active viral replication relies on the detection of a replicative intermediate, the negative ssRNA strand [25, 34]. However, these non-encapsidated RNA molecules are unprotected and prone to rapid degradation following cell death. Virions of several viral lineages undergo various degrees of maturation before becoming infectious [35–37]. Although these changes are subtle in picorna-like viruses [38], we could still take advantage of the structural characterization of immature particles, as indicators of active viral replication. We demonstrated this at near-atomic resolution of only 312 (2.57 Å) and 832 (2.35 Å) particles by detecting the uncleaved VP0, a hallmark of picorna-like provirions [39–41]. As VP0 cleavage is irreversible, the presence of uncleaved VP0 cannot result from postmortem degradation of mature virions, supporting the identification of authentic provirion populations. This process only required 20 honey bee carcasses, which were deceased for several days prior to virus purification.

Our diagnostic process also produced the structural characterization of the ABPV mature, vectored virion and its two immature forms. The interpretation of the fivefold-related external density, a unique ABPV feature, is challenging, as the residues involved in its coordination are all conserved in KBV and IAPV. Additionally, both the IAPV and KBV reconstructions are based on honey bee-derived particles, which argues against a general host-specific origin of this density [19–21]. The dynamics of the VP2(VP0) C-termini are also unique to ABPV, as KBV and IAPV exhibit fully ordered VP2 C-termini. This absence exposes the base of the surface spikes, which could be necessary to allow mature virions to bind specific receptors. ABPV virions are transported to the hypopharyngeal- and the salivary glands of infected honey bees following their egress to the hemolymph, which are both rich in proteases [42]. Consequently, the C-termini may be removed by cleavage, rather than failing icosahedral averaging.

The characterization of ABPV provirions provided structural evidence of viral replication and assembly in Colony B, which exhibited paralytic signs. This also provided structural insights into the maturation process of aparaviruses, which is consistent with a maturation model, where initially the ssRNA genome is in a compact, intrinsically ordered form, which gradually expands to occupy the capsid lumen. The significant void between the genome and the capsid interior provides sufficient space for the VP0 N-termini to move around freely, manifesting in its lack of icosahedral ordering. As the genome expands and the void decreases, its movement is restricted, which manifests in its increasing tendency to follow the icosahedral order of the virion shell. The cleavage of VP0 to VP2 and VP4 may be the consequence of this restricted movement, stabilizing the interaction between the VP0 cleavage site and the DDF autocatalytic motif of VP3. The expansion of the genome might be attributed to forces exerted by positively charged surfaces of the capsid interior, occupying the fivefold- and threefold axes, as well as the VP4 C- and N-termini (Fig. S6). Members of the *Iflaviridae*, another picorna-like virus family, which includes the honey bee-infecting deformed wing virus, were found to package their genomes into pre-formed empty capsids [18]. As empty, pre-assembled particles were absent from the immature fractions, yet the provirions exhibited a compact genomic core, our findings suggest that ABPV, and possibly other dicistoviruses, exhibit a fundamentally different maturation mechanism from that of iflaviruses and assemble their capsids around an intrinsically more ordered, compact genomic core.

Despite sporadic PCR-based detections of ssRNA from honey bee picorna-like viruses, the role of the SHB as a virus vector is little understood [34, 43]. Our findings imply that SHBs carry a significant number of intact, mature ABPV virions. None of the four independent colonies exhibited ABPV-negative beetles. ABPV particles were absent from the environment, i.e. beeswax, in Colony C despite the high SHB larval titers, implying that the larvae may have contracted the virus from a source outside of the collapsed colony. As adult beetles lay eggs directly into the hives, a vertical mechanical transmission from adult carriers could be plausible, given the demonstrated environmental stability of ABPV. Intact, mature ABPV virions were present both on the cuticle and in the bodies of SHB larvae, following colony loss, even several days later. Although immature virions could not be detected in the SHB, this observation does not warrant their complete exclusion from being potential biological vectors, which requires further evaluation.

## Conclusions

Using honey bee colony losses as a diagnostic challenge, we could demonstrate the capabilities of a structure-first approach even on degraded environmental samples and carcasses, using only a limited amount of material and viral particles. Our case study revealed distinct aparaviral maturation states consistent with a model, in which genome reorganization precedes VP0 cleavage. As SHBs demonstrated a strong association with ABPV in these lost colonies, their role in introducing viral agents to healthy colonies may pose an additional risk beyond their destructive presence. Cryo-EM, as a diagnostic tool, has the capability to identify a viral pathogen, assess its active replication status, and reveal insights into the viral structural biology, directly from field-collected specimens—without requiring prior sequence-based information or cell cultures. This combination of diagnostic and biological information supports the further development of structure-first approaches for investigating agricultural disease outbreaks.

## Material and Methods

### Origin of the samples

Samples of independently lost colonies were obtained from beekeepers located in Bastrop (colony A), Guadalupe (colony B), and Victoria (colony C) in southeastern Texas. The remains of each colony were collected by the beekeepers and were either shipped directly to our laboratory or were transported by TAIS. Colony A showed a gradual die-off of bees following the increased presence of SHBs. Eventually, surviving workers either abandoned the hive or failed to navigate back to it, leaving an empty beehive completely infested with SHB larvae. Colony B showed gradual die-off of both workers and brood in a one-week time frame. Colony C showed a sudden surge in SHB population, accompanied by an increase in honey bee worker deaths. All three colonies exhibited prominent SHB infestation. Colony D, a healthy colony of TAIS, located in Bryan, Texas, and served as a healthy control.

### Purification of viral particles

For Colony A, hive remains of fragmented comb, beeswax, and SHB larvae were mechanically homogenized in 1×PBS pH 7.2 (137 mM NaCl, 2.7 mM KCl, 10 mM Na_2_HPO4, 1.8 mM KH_2_PO_4_, pH). The homogenate was combined with an equal volume of 1×TNTM pH 8 (50 mM Tris at pH 8, 100 mM NaCl, 0.2% Triton X-100, 2 mM MgCl_2_) and the lysate was clarified by repeated centrifugation at 4,300 × g for 15 min at 4 °C. The supernatant was mixed with 1×TNET pH 8 (50 mM Tris pH8, 100 mM NaCl, 0.2% Triton X-100, 1 mM EDTA) in a 1:3 ratio and concentrated on a cushion of 20% w/v sucrose in TNET using a type 45Ti Beckman-Coulter ultracentrifuge rotor for 3 h at 10 °C at 42,300 rpm on a Beckman Coulter S-class ultracentrifuge. The pellet was resuspended in 1 mL of 1×TNTM pH 8 and, following overnight incubation, it was purified on a 5-70% w/v sucrose step gradient of 5% increments. The gradient was spun at 10°C for 3 h at 35000 rpm, using the same S-class instrument with a SW41Ti swinging bucket rotor.

The positive fractions scattered the light only faintly, and we pulled all fractions exhibiting the slightest blue color. These fractions were aspirated by a single needle puncture and a 10-mL volume syringe. The purified fraction was dialyzed against 1×PBS, pH 7.4 to remove the sucrose. The dialyzed sample of ∼5 mL volume was concentrated 10-fold on an Amicon^®^ Ultra centrifugal filter unit of 100 kDa molecular weight cut-off.

From Colony B, we randomly selected 20 honey bee carcasses, which were homogenized in 30 mL 1×TDX, pH 7.4 (137 mM NaCl, 2.7 mM KCl, 10 mM Na_2_HPO_4_, 1.8 mM KH_2_PO_4_, 1 mM MgCl_2,_ 0.2% Triton X-100), using a glass mortar. The homogenate was processed through the same subsequent steps as the remains of Colony A.

### Grid preparation, vitrification, and cryo-transmission electron microscopy

For the detection and identification of ABPV, Quantifoil R1.2/1.3 300-mesh copper grids with a 2-nm-thick, commercially generated continuous carbon film were glow discharged on an easiGlow™ system. The purified immature ABPV virions were vitrified on UltrAufoil R1.2/1.3 300-mesh gold cryo-EM grids. Samples were plunge-frozen into liquid ethane using a ThermoFisher Vitrobot Mark IV at 100% humidity and ambient temperature after blotting away the excess liquid for 5 s at 0 force. The grids were clipped into autoloader grids and imaged using a ThermoFisher Titan Krios G4 Cryo-transmission electron microscope with a K3-direct electron detector camera, operated in low dose mode at 300 kV, with a slit width of 20 eV or 15 eV for the two high resolution data collections, respectively. Images were recorded with a 100 μm objective aperture, at a 2.25-s-long exposure, with a total dose of ∼50 e^-^/Å^2^, using a frame length 0.2 s for both collections. Movie frames were recorded in counting mode using the ThermoFisher EPU software [44] at a sampling of 1.068 Å/pixel (Colony A) and 0.832 Å/pixel (Colony B).

### Cryo-EM data processing and atomic model building

The collected movies were processed in the cryoSPARC 4 suite [45]. Movie frames were aligned by the Patch Motion Correction subroutine, and the quality of the aligned micrographs was assessed by the Patch CTF Estimation subroutine. Boxed particles were extracted and subjected to 2-dimensional classification. Particles of classes that failed to display a clear 2D-class average of the icosahedral particle were eliminated from the reconstructions. The particles from Colony B were further classified by reference-free 3-dimensional classification, using the “simple” classification option. All 3D classes underwent individual processing. *Ab initio* reconstruction, due to the low number or particles, was performed at a 12 Å initial alignment resolution with incorporating all particle images into the estimation of the signal-to-noise prior, imposing icosahedral symmetry. The obtained startup volume was subjected to automatic refinement under icosahedral constraints and underwent iterations until pseudoconvergence. To improve the resolution, corrections for higher-order aberrations, beam tilt, trefoil and anisotropic magnification were applied, as well as per-micrograph astigmatism and per-particle CTF parameters. The automatic high-resolution refinement was repeated in the presence of a soft-edged mask, using the Non-Uniform Refinement subroutine. The resolution of each reconstructed map was calculated based on a Fourier shell correlation (FSC) of 0.143.

The cryo-EM maps were subjected to both *de novo* and sequence-specific model building using ModelAngelo [22]. As automated modeling of VP4 was unsuccessful, the corresponding density was identified manually. VP4 was visible only at higher map-contour levels and was manually traced in Coot [46]. The raw atomic model of all four VPs was refined into the density using ISOLDE [47], followed by further refinement cycles in Coot, and other subsequent refinement cycles in ISOLDE. The best refined atomic model was then subjected to automated refinement in PHENIX [48], in the presence of NCS restraints through five macrocycles and an eventual B-factor refinement. Visualization and the preparation of publication quality figures was performed using UCSF Chimera[49] and ChimeraX [50].

### Metagenome sequencing and assembly

Viral RNA was extracted from 50 uL purified virus suspension, using the Qiagen RNeasy kit. Extracted RNA was subjected to reverse transcription using the SuperScript III reverse transcriptase (Thermo Fisher Scientific) in the presence of random hexamers. The reverse-transcribed cDNA was shipped to a commercial second-strand synthesis service followed by Illumina sequencing provided by Celemics (Seoul, Korea). The raw Illumina reads were trimmed using Trimmomatic v0.39 [51]. The evaluation and filtering of read quality was conducted by Fastp v0.23.4 [52], and *de novo* assembly was carried out by MEGAHIT v1.2.9 [53]. Assembled contigs were screened using geNomad v1.12.0 [54] to identify and classify viral contigs.

### Phylogenetic and structural comparisons

The asymmetric unit model (ASU) of ABPV was compared against that of IAPV (PDB accession: 5LWG) and KBV (PDB accession: 7BGK) using matchmaker on UCSF Chimera to calculate Root-Mean-Square Deviations (RMSD) with the Needleman-Wunsch algorithm with the BLOSUM-62 residue similarity matrix (weight 0.7) and secondary structure scoring (weight 0.3). RMSD was calculated for each VP separately. The amino acid sequence of each VP of ABPV (Genbank: PZ621326 and Q9DSN8), IAPV (PDB: 5LWG), KBV (PDB: 7BGK) and the outgroup Solenopsis invicta virus (SINV) (GenBank: AAU85376) were aligned using MAFFT [55]. Phylogenetic inferences were calculated using RaxML [56] within the Geneious Prime suite (https://www.geneious.com/).

### PCR screenings and RNA quantification

For reverse transcription the primer 5’-TCTGGAACTGCAGAAGATAAAGCT-3’ was used, targeting the VP1-coding region of the genome. Total RNA of small hive beetles, wax, and honey bee samples were extracted using a TRIzol-chloroform protocol. Briefly, each extraction included a single deceased insect or 40 mg beeswax, all of which were homogenized using a sterile pestle in a sterile microcentrifuge tube. 1 mL TRIzol was added to each homogenate. Insect samples were incubated for 15 min, while beeswax pieces were subjected to 1.5-h-long incubations, which proved to be sufficient to completely dissolve the wax. Following vigorous mixing, the samples were incubated for 5 min, followed by pelleting at 12000×g for 15 min. The aqueous phase was subjected to RNA precipitation by adding 500 µL isopropanol. The precipitated nucleic acid was pelleted at 12000×g for 10 min at 4°C, following a 10-min-long incubation. The RNA pellet was washed in 1 mL ice cold ethanol, and resuspended in 20 µL nuclease-free water once the ethanol evaporated. The purified RNA was subjected to the previously described reverse transcription protocol.

For PCR screening, we designed a second primer pair of 5’-CAGCCCCAGTGAATCCTCTA-3’ and 5’-ACTACAGAAGGCAATGTCCAAGA-3’ targeting an approximately 400-nt-long region nested within the cDNA products. Amplifications were executed in a reaction volume of 25 μL, including 2 μL of cDNA target, 0.5 μL of both primers in 50 pmol concentration, 0.5 μL dNTP mix with 8 μmol of each nucleotide, and 0.1 μL of DreamTaq DNA polymerase enzyme (Thermo Fisher). PCR reactions were executed under a program of 5 min denaturation at 95°C followed by 35 cycles of 30 s denaturation at 95°C, 30 s annealing at 50°C, and 1 min of elongation at 72°C, and a final elongation step of 5 min at 72°C. The resulting products were validated using a commercial Oxford nanopore amplicon sequencing service, provided by Plasmidsaurus (Eugene, OR, USA).

Quantification of the viral RNA was carried out by qRT-PCR amplification in a BioRad CFX Duet instrument. A 400-nt-long target sequence was amplified and quantified by the same primer pair designed for the PCR screenings, using the BioRad iTaq™ Universal SYBR^®^ Green Supermix with an amplification program of 5-min denaturation at 95 °C followed by 45 cycles of 30 s denaturation at 95°C, 15 s annealing at 55°C, and 30 s of elongation at 72°C. Results were analyzed with the CFX Maestro Software (BioRad). All reactions were carried out in triplicates for all biological samples in all experimental assays. To establish the accurate viral titer and to avoid solely relying on Cq values, we cloned the 1124-nt-long amplification product of the primer pair used for reverse transcription into a pJET1.2/blunt cloning vector, the concentration of which could be measured and confirmed by a series of measurements in a spectrophotometer. This plasmid was used as the target DNA of known concentration to calculate the standard curve for each quantification run.

### Limit of detection assay

To determine the detection threshold of the PCR and qPCR screenings, a limit of detection assay was performed using the purified and concentrated ABPV virus suspension of Colony A. This stock was used to create a 10-fold dilution series from 1 to 10^-10^. RNA was extracted from each element of the series and quantified as three technical replicates.

To compare the quantification results to the amount of viral genomes used in the cryo-EM grid preparation process, two identical Quantifoil 1.2/1.3 300-mesh grids with a continuous carbon film were glow discharged and blotted for 5 seconds at a blot force 0 using a ThermoFisher Vitrobot Mark IV with the original ABPV suspension. After soaking and washing each grid in 1×PBS, supplemented with 0.2% Triton X-100, the RNA was extracted from each wash separately, and the viral genome copies were quantified using the qRT-PCR assay.

### Negative staining electron microscopy and quantification of the internal and external SHB-associated ABPV titer

Three larval carcasses were subjected to vigorous wash in 1 mL 1× sterile PBS, supplemented with 0.2% Triton X-100. The post-wash buffer was collected and concentrated to a final volume of 50 µL for RNA extraction and viral titer quantification. The washed larvae were processed separately for RNA extraction and quantification of viral titers within larval tissues. The homogenate was cleared by centrifuging for the negTEM experiments. A 3 µL aliquot of the cleared homogenate and the PBS suspension was applied to previously glow-discharged TEM grids with a thin carbon film (MiTeGen) and was stained by 1% uranyl acetate. The grids were imaged by an FEI transmission electron microscope at a 40 kV acceleration voltage.

### Viral titer quantification of hive-related materials

Viral loads were quantified across three hive-associated samples from each colony, including SHB larvae or adults, beeswax from combs, and carcasses of honey bees. For each sample, RNA was extracted separately from eight biological replicates per colony, and viral loads were quantified by qRT-PCR using three technical replicates per sample, following reverse transcription.

### Time course persistence of viral particles in beeswax

Beeswax wax collected from fragmented combs of Colony A, which was previously stored at −20°C following its reception at our laboratory. The beeswax fragments were incubated at ∼30°C sampled at four time points of a week-long incubation interval on day 0, day 1, day 3, and day 7. At each time point, RNA was extracted from three biological replicates, followed by qRT-PCR quantification of the ABPV titer. Each biological replicate was quantified using three technical replicates. Reactions with Cq values >35 were considered as low-confidence detections near the detection limit of the assay.

### Differential Scanning Fluorometry (DSF) assay

Purified ABPV particles at 0.05 mg/mL concentration in 1× PBS, pH 7.2, were mixed with either 1% SYPRO^TM^orange dye (Invitrogen) or with 0.5% SYBR^TM^Gold (Invitrogen) dye. The stained particles were incubated in a BioRad CFX Duet Real-Time PCR instrument with active detection at the FRET and SYBR detection channels. From 30°C to 99°C, the specimen was screened at a ramp rate of 1 °C/min in 0.5 °C increments. Fluorescence was measured as the function of temperature, plotted as -dRFU/dT vs. temperature, which was multiplied by −1 and normalized to the highest RFU value of each well.

## Acknowledgements

We thank Angela Steinhauer and Hannah Blackburn (Texas Apiary Inspection Service), Dr. Tonya Shepherd and Dr. Juliana Rangel (Texas A&M Honey Bee Lab), and the Texas beekeepers who provided the sample material used in this study. We thank Dr. Joseph Szule for his assistance on negTEM data collection at the Image Analysis Laboratory at Texas A&M University. We thank Dr. Gaya P. Yadav for cryoEM data collection at the Laboratory for Biomolecular Structure and Dynamics (LBSD) of Texas A&M University. The LBSD is supported, in part, by the Department of Biochemistry & Biophysics, AgriLife, Texas A&M University, and CPRIT Facility Support Grant (RP25049). Funding for this study was provided by the Texas A&M AgriLife Insect Vector Program, Grant No. 06-124196-81240, awarded to PI Penzes.

## Data availability

Nucleic acid sequence data generated in this study has been deposited to the NCBI GenBank under accession number PZ62132. Density maps generated via cryo-electron microscopy were deposited to the Electron Microscopy Data Bank under accession numbers of EMD-78018, EMD-78025, and EMD-78035. The atomic models corresponding to each map were deposited to the RCSB Protein Data Bank, under accession numbers 37BJ, 37AX and 37AZ. The raw cryo-EM datasets generated and analyzed in this study are available from the corresponding author (J.J.P.) (by mail at) upon reasonable request as part of a scientific collaboration with adherence to standards of good scientific practice, although restrictions may apply due to privacy reasons and ongoing research projects. The computational codes used to analyze raw qRT-PCR data are available at https://github.com/jorgemedinad/.

